# Exogenous L6 myotube mitochondrial transplantation attenuates hypertrophy and elicits a unique proteomic signature in phenylephrine-treated H9C2 cardiomyocytes

**DOI:** 10.64898/2026.09.21.753229

**Authors:** Nicholas J. Kontos, George J. Kontos, Dustyn T. Lewis, Samuel C. Norton, Bradley A. Ruple, Andreas N. Kavazis, Darren T. Beck, Melissa D. Boersma, Christopher B. Mobley, Danielle J. McCullough, Ahmed Ismaeel, Michael D. Roberts

## Abstract

Mitochondrial transplantation has recently emerged as an alternative treatment for cardiovascular disease (CVD), aimed at increasing mitochondrial number and improving mitochondrial function. Though mitochondrial dysfunction is a key factor in the development of right ventricular hypertrophy/failure, the effects of mitochondrial transplantation on disease mitigation have been largely unexplored. Therefore, this *in vitro* study aimed to determine whether the transplantation of exogenous L6 myotube mitochondria mitigates negative outcomes in H9C2 cardiomyocytes that were stimulated to hypertrophy with phenylephrine. Control (CTL), control with mitochondrial transplantation (MitoTx), Phenylephrine only (Phe-only), and phenylephrine with mitochondrial transplantation (Phe+MitoTx) treatments were evaluated. Pilot experiments indicated mitochondrial transplantation into healthy cardiomyocytes acutely increased Complex I-linked oxidative phosphorylation (OXPHOS) capacity (p=0.005) and maximal respiratory capacity (p=0.022) within 24 hours, and these data alongside microscopic evidence of fluorescently labeled L6 mitochondria in H9C2 cardiomyocytes suggested successful transplantation. Regarding treatment comparisons, Phe-only showed a significant increase in cell area (p<0.05), while Phe+MitoTx blunted the hypertrophic cardiomyocyte response. Consistent with these results, proteomic analysis of 4,806 proteins showed that transplantation enriched the mitochondrial proteome, impacting pathways including OXPHOS, respiration, fatty acid and amino acid metabolism, while suppressing extracellular matrix remodeling and de-differentiation signatures. This response was observed in healthy and phenylephrine-stressed cardiomyocytes. In conclusion, our *in vitro* data indicates that transplantation of L6 skeletal muscle mitochondria mitigates negative effects induced by phenylephrine in H9C2 cardiomyocytes. However, more rigorous *in vivo* studies are needed to determine if this is a suitable approach for disease mitigation.

## INTRODUCTION

Right ventricular (RV) failure is characterized by the inability to effectively pump deoxygenated blood into the lungs, and this disease carries a poor prognosis and frequently arises secondary to pulmonary hypertension, ischemic heart disease, or congenital heart defects (1, 2). RV failure features compensatory hypertrophy, progressive dilation, and decompensation. Unlike the left ventricle, the thin-walled RV is optimized for high-volume, low-pressure work, making it poorly equipped to tolerate acute pressure overload imposed by pulmonary disease, which precipitates a dramatic drop in cardiac output (3). The result is elevated right-sided pressures, systemic venous congestion, and significant morbidity and mortality (4).

Central to this deterioration is a failure of cardiac energetics. Notably, the heart is the body’s most metabolically active organ, requiring continuous ATP synthesis to sustain its function. In a healthy/normoxic heart, over 95% of cardiac ATP is generated via mitochondrial oxidative phosphorylation (OXPHOS), with the remainder derived from anaerobic glycolysis (5). In addition, this cardiac OXPHOS is primarily driven by oxidation of fatty acids (70-90%), with the remaining coming from the oxidation of glucose and lactate (10-30%) (5). RV failure is associated with mitochondrial dysfunction, resulting in reduced energy yield, increased reactive oxygen species (ROS) production, and inefficient substrate metabolism (6). This results in decreased oxidative phosphorylation activity due to increased pyruvate dehydrogenase kinase activity, which prevents pyruvate from entering the Krebs Cycle (7). The heart transitions from fatty acid oxidation to glucose utilization via anaerobic glycolysis for ATP production (8), compounding mitochondrial dysfunction and further depressing RV contractility (7).

Though progress has been made in treating RV failure, much of the therapeutic focus has been on mitigating fluid overload (9). Conversely, therapeutics related to cellular resilience and bioenergetic restoration have largely been overlooked. Although pharmacologic manipulation of mitochondria alone may modulate defective energy production, it is not sufficient to treat the failing right ventricle (10). Mitochondrial transplantation has recently emerged as an alternative treatment for cardiovascular disease, aimed at increasing mitochondrial number and improving mitochondrial function (11). Preclinical studies have demonstrated that transplantation of viable, respiration-competent mitochondria into ischemic cardiac tissue can restore ATP production, reduce oxidative stress, and attenuate cardiomyocyte apoptosis, thereby preserving contractile function (12–14). These beneficial effects have been largely characterized in the context of left ventricular ischemia-reperfusion injury, where mitochondrial dysfunction is a well-established driver of pathology. However, limited evidence exists regarding the effects of mitochondrial transplantation on RV failure (15).

Therefore, we employed an *in vitro* model to determine whether the exogenous transplantation of myotube mitochondria mitigates outcomes associated with RV hypertrophy and dysfunctional bioenergetics in H9C2 cardiomyocytes stimulated with phenylephrine. The experiments performed herein are described as AIM 1 and AIM 2. The chief purpose of AIM 1 was to establish and optimize methods leading to the successful uptake of exogenous mitochondria in cardiomyocytes. The purpose of AIM 2 was to determine whether exogenous mitochondrial transplantation improves cardiomyocyte bioenergetics and attenuates the hypertrophic and proteomic changes induced by phenylephrine. We hypothesized that transplantation would improve bioenergetics, coinciding with reductions in cellular hypertrophy and a partial reversal of the phenylephrine-induced proteomic signature.

## METHODS

### AIM 1 Mitochondrial isolation from L6 myotubes

#### L6 culture

L6 immortalized rat myoblasts (ATCC, Manassas, VA, USA), with passages 1-3, were cultured at 37°C in a 5% CO_2_ atmosphere. Myoblasts were seeded at a density of 2.2 × 10^6^ in 100 mm plates containing 10 mL of growth media (GM) consisting of Dulbecco’s modified eagle’s medium (DMEM; Corning, Corning, NY, USA), 10% Fetal Bovine Serum (Corning), 1% penicillin/streptomycin (VWR, Radnor, PA, USA), and 0.1% gentamycin (VWR). Upon reaching confluency (∼80–90%), myoblasts were differentiated into myotubes by switching to differentiation media (DM) containing DMEM, 2% horse serum (VWR), 1% penicillin/streptomycin (VWR), and 0.1% gentamycin (VWR). DM was replaced every 24–48 h until mature myotubes reached confluence (∼7 days). Following 7 days of differentiation, myotubes were washed twice with sterile phosphate buffered saline (PBS; Corning) and then incubated with 0.25% Trypsin for 5 minutes (Corning). Cells were pelleted by gentle centrifugation (∼300×g, 5 min) and were then subjected to mitochondrial isolation using the Mitochondrial Isolation (MitoIso) Kit for Cultured Cells (Thermo Fisher Scientific, Rockford, IL, USA; cat. #: 89874). The isolation steps that follow were performed on ice to preserve the mitochondria.

#### Mitochondrial isolation from L6 myotubes

Protease inhibitors provided by the kit were added to MitoIso Kit Reagent A and Reagent C. The supernatant of the cell pellet contained in a 2.0 ml microcentrifuge tube was carefully removed and discarded. Then, the pellet was dissolved in 800 µL of MitoIso Reagent A, vortexed on a benchtop device at medium speed for 5 seconds and incubated on ice for exactly 2 minutes. The cell suspension was then transferred into a prechilled Dounce Tissue Grinder. The cells were lysed by performing 70 strokes in the grinder on ice. The lysed cells were returned to the original tube and 800 µL of MitoIso Reagent C was added. The Dounce Tissue Grinder was rinsed with 200 µL of Reagent A, which was then added to the sample tube. The tube was inverted several times to mix. Then, the tube was centrifuged at 700 × g for 10 minutes at 4°C. The supernatant was transferred to a new 2.0 ml tube and centrifuged at 12,000 x g for 15 minutes at 4°C. The resultant supernatant (cytosol fraction) was transferred to a new tube. The pellet, containing isolated mitochondria, was washed with 500 µL Reagent C and centrifuged at 12,000 x g for 5 minutes at 4°C, and the supernatant was carefully removed and discarded. The mitochondrial pellet was kept on ice before downstream processing, as we operated under the premise that freezing and thawing would compromise mitochondrial integrity.

#### L6 mitochondrial isolate protein quantification

The mitochondrial pellet protein analysis was performed using a commercially available BCA protein assay kit (Thermo Scientific, Waltham, MA, USA; cat. #: A55864) and spectrophotometer (Agilent Biotek Synergy H1 hybrid reader; Agilent, Santa Clara, CA, USA). For protein analysis, the mitochondrial pellet was dissolved in 100 µL of 1x general cell lysis buffer (Cell Signaling) and vortexed for 1 minute. The mitochondrial lysate was centrifuged at 500 x g for 5 minutes at 4°C. The supernatant containing the soluble mitochondrial protein was analyzed by the BCA protein assay microplate procedure according to the manufacturer’s instructions. Notably, 1x general cell lysis buffer was used to generate the serial BCA standard curve instead of deionized water to account for potential background effects.

### AIM 1 mitochondrial transplantation of L6 myotube mitochondria into healthy H9C2 cardiomyocytes

H9C2 cells were originally derived from embryonic rat heart tissue by Kimes and Brandt in 1976 (16), and were obtained from a commercial vendor (ATCC, Manassas, VA, USA). Passages 1-2 cells were cultured at 37°C in a 5% CO_2_ atmosphere and seeded at a density of ∼3 × 10^5^ in six-well plates containing 3 mL/well of growth media (GM) consisting of DMEM, 10% Fetal Bovine Serum (Corning), 1% penicillin/streptomycin (VWR), and 0.1% gentamycin (VWR). When cells reached ∼80–90% confluency, the L6 myotube mitochondrial pellet (generated as described in the preceding paragraphs under mitochondrial isolation from L6 myotubes) was dissolved in room temperature GM, and the resuspended mitochondria were added to each well and co-incubated with cardiomyocytes for a period of 24 hours at 37°C in a 5% CO_2_ atmosphere.

#### Fluorescent labeling and imaging of L6 mitochondrial transplantation

For certain AIM 1 and AIM 2 experiments, L6 myotubes were incubated with MitoTracker Red FM (Thermo Scientific, cat. #: M46751) (ex/em 581/644 nm) for 45 minutes at 37°C (protected from light) prior to the MitoIso procedure and washed twice with PBS to remove unbound dye. These mitochondria were then incubated with H9C2 cardiomyocytes that were seeded on CultureWell MultiWell chambered coverslips (AIM 1) (Thermo Fisher Scientific; cat #: C24776) at 50,000 cells per well 24 hours prior to mitochondrial transplantation or fibronectin-coated glass coverslip discs (AIM 2) (Corning BioCoat, cat #: 354008) at 300,000 cells per well 24 hours prior to PE treatment. Various doses of L6 mitochondrial isolates were piloted (AIM 1). On a given 2-well chambered coverslip, the first well was labeled ‘control’ and contained no exogenous L6 mitochondria (Vehicle PBS). A low dose (1.25 µg of protein according to BCA readings), a moderate dose (2.5 µg protein), and a high dose (5 µg protein) of L6 mitochondrial isolates were applied to other wells. After 24 hours of co-incubation at 37°C/5 % CO_2,_ the GM was aspirated, each well was washed with PBS, the gasket was removed, and a glass coverslip was applied to the slide. Care was taken to ensure that the GM from each well did not evaporate during the co-incubation period in the incubator. Multiple images were obtained with a fluorescent microscope using a 20× objective (Zeiss Axio imager.M2; Zeiss). Imaging was used to confirm internalization of exogenous mitochondria into cardiomyocytes.

#### High Resolution Respirometry

For certain AIM 1 and AIM 2 experiments, cardiomyocytes (∼1 × 10^6^ cells) were harvested, following 24 hours of co-incubation (with mitochondria or vehicle) (AIM 1) or phenylephrine treatment (AIM 2), and added to respiration media consisting of Mir05 buffer supplemented with 20 mM creatine monohydrate (Sigma Aldrich; Cat #: 27900). A substrate inhibitor titration (SUIT) protocol was performed to assess complex-specific mitochondrial respiration using an Oroboros Oxygraph-2 k (O2k) FluoRespirometer (Oroboros Instruments, Innsbruck, Austria). After permeabilization with digitonin (0.02mg/mL) (Sigma Aldrich; cat. # D5628), pyruvate (5 mM) (Sigma Aldrich; cat #: P2256) and malate (2 mM) (Sigma Aldrich; cat #: M1000) were added to the oxygraphy chambers to measure basal complex I, state 2 (LEAK) respiration. This was followed by ADP (4 mM) (Sigma Aldrich; cat. # 117105) addition to initiate state 3 (Complex I-linked OXPHOS) (ADP-stimulated) respiration. Succinate (10 mM) (Sigma Aldrich; cat. #: S2378) was added to stimulate electron flow through Complex II, followed by a titration of carbonyl cyanide m-chlorophenyl hydrazine (CCCP, 0.25 μM to 1.5 μM) (Sigma Aldrich; cat #: C2759) to stimulate maximal uncoupled respiration (ET capacity). Rotenone (10 μM) (Sigma Aldrich; cat. #: R8875) was used to inhibit complex I, and 5 μM antimycin A (Sigma Aldrich; cat. #: A8674) was used to inhibit electron flow through complex III, accounting for non-mitochondrial oxygen consumption. Finally, Complex IV-linked OXPHOS was determined using 0.4 mM N,N,N’,N’-tetramethyl-p-phenylenediamin (TMPD) (Sigma Aldrich; cat. #: T3134) and 2 mM ascorbate (Sigma Aldrich; cat. #: A7631) to prevent TMPD auto-oxidation. Cytochrome c addition (10 μM) (Sigma Aldrich; cat. #: C7752) was used to assess potential damage to the outer mitochondrial membrane, with increments of oxygen flux response after cytochrome c addition below 15%. After completion of the SUIT protocol, samples were removed from the oxygraphy chambers, cell count was measured using a hematocytometer, and the respiration rate was normalized to cell count.

### AIM 2 assessments of L6 myotube mitochondrial transplantation effects on H9C2 cardiomyocytes treated with phenylephrine

H9C2 cardiomyocytes (passage numbers 2-3) were cultured at 37°C in a 5% CO_2_ atmosphere. Cardiomyocytes were seeded at a density of ∼3 × 10^5^ in six-well plates containing 3 mL/well of growth media (GM) consisting of DMEM, 10% Fetal Bovine Serum (Corning), 1% penicillin/streptomycin (VWR), and 0.1% gentamycin (VWR). When the cardiomyocytes reached ∼80–90% confluency, cells were treated with low-serum DMEM containing 1% FBS (Corning), 1% penicillin/streptomycin (VWR), and 0.1% gentamycin (VWR) for 18 hours. Four independent treatments were then applied to 6-well plates for 48 hours including: i) control (CTL), whereby low serum DMEM was applied to cells two times over a 48-hour period (24 hours per), ii) control with mitochondrial transplantation (CTL+MitoTx), whereby low serum DMEM was applied to the cells two times over a 48 hour period (24 hours per) as well as isolated L6 Mitochondria (∼5 ug protein) was applied to cells for the second 24 hour period, iii) phenylephrine-only (Phe-only), whereby low serum DMEM with 100 µM phenylephrine (Thermo Fisher Scientific; cat. #: 20724050) was applied to cells two times over a 48-hour period (24 hours per) (17), or iv) phenylephrine with mitochondrial transplant (Phe+MitoTx), whereby low serum DMEM with 100 µM phenylephrine was applied to cells over a 48-hour period (24 hours per) as well as isolated L6 mitochondria (5 µg protein) was applied to cells for the following 24 hours, with the low serum DMEM + 100 µM phenylephrine present in the media. Following this treatment scheme, experiments included: i) cells being fluorescently labelled on 6-well plates using a phalloidin-conjugated Alexa Flour 488/DAPI protocol for cell size imaging with concomitant fluorescent labeling of internalized L6 mitochondria (treatments i, iii, iv above), b) high resolution microscopy as described in AIM 1 (treatment i and iii), or c) lysed using general cell lysis buffer and subjected to proteomics (described in a later section).

#### Phalloidin Stain for Cell Area and Quantification

Following coincubations described in the preceding section, cells were washed twice with PBS and incubated for 10 minutes at room temperature in 4% paraformaldehyde (PFA). Following fixation, cells were washed three times with PBS. After siphoning off the last PBS wash, cells were permeabilized with 0.1%Triton X-100 (Amresco, Solon, OH, USA; cat. # 0694-1L) in PBS for 10 minutes at room temperature. After washing three times with PBS, filamentous actin (F-actin) was stained by incubating cells with green-fluorescent Alexa Fluor 488 Phalloidin (Invitrogen, Carlsbad, CA, USA; cat. # A12379) diluted 1:400 in PBS for 1 hour at room temperature, protected from light. Following three washes with PBS, cells were incubated with DAPI (1:10,000) (Thermo Fisher Scientific, Waltham, MA, USA; cat. #: D3571) at room temperature for 10 minutes. Cells were then washed and moved to the microscope for imaging. Digital images were captured using a fluorescent microscope (10x objective; Zeiss Axio imager.M2) and motorized scanning stage. Fluorescence images were quantified using ImageJ software (National Institutes of Health; Bethesda, MD, USA). For experimental rigor, three treatment groups were quantified across three separate wells per group, with five random images captured per well. Again, digital images were analyzed using ImageJ software. Spatial scale calibration was established using the “Set Scale” function. To avoid sampling bias, all intact H9C2 cells located entirely within the field of view with clearly discernible boundaries were manually outlined using the freehand selection tool, yielding approximately 20-30 analyzed cells per image. Cell areas were averaged to calculate the final total cell area per well.

#### Proteomic Analysis and Bioinformatics

Proteomics were performed on the lysates similar to previous work published by our laboratory (18). Treated cells were studied under a balanced 2×2 design crossing phenylephrine-induced hypertrophic stress with mitochondrial transplantation (four groups CTL, MitoTx, Phe-only, Phe+MitoTx; six replicates each, 24 samples). Three contrasts were performed for simple effects including “Disease” (Phe-only − CTL), “Transplant” (MitoTx − CTL), and “Rescue” (Phe+mitoTx – Phe-only). The fourth contrast represented their interaction ((Phe+MitoTx − Phe-only) − (MitoTx − CTL)). Transplant and Rescue draw on separate groups, so they were statistically independent. Disease and Rescue both involve the Phe-only group and are therefore correlated by construction (r = −0.50); comparisons that span this pair are judged against a label-shuffling permutation null (300 permutations, seed 42). At six replicates per group, the Interaction is underpowered. Two secondary contrasts, reported in S1 Table, complete the design: Disease-after-transplant (Phe+MitoTx − MitoTx), the phenylephrine effect in transplanted cells, and Recovery (Phe+MitoTx − CTL), the net shift of the rescued state from control.

Protein lysates (50 µg per sample) were prepared with the EasyPep Mini MS Sample Prep Kit (Thermo Fisher; cat. #: A4006), with reduction and alkylation at 95°C for 10 min, Trypsin/Lys-C digestion at 37°C for 2 h, and peptide clean-up per the kit instructions. Peptides were analyzed on an externally calibrated Thermo Orbitrap Exploris 240 coupled to a Dionex UltiMate 3000 RSLC Nano system (Thermo Fisher): samples were loaded onto an Acclaim PepMap 100 trap column (75 µm i.d. × 2 cm) and separated at 300 nL/min on an Easy-Spray PepMap RSLC C18 analytical column (50 µm × 15 cm) over a 135-min linear gradient from 3% to 50% B (mobile phase A, 99.9% H₂O / 0.1% formic acid; B, 80% acetonitrile / 0.1% formic acid), with direct nanospray into the instrument under Thermo Xcalibur control. MS1 spectra were acquired from 380–985 m/z at 60,000 resolution. MS2 spectra were acquired in data-independent mode with a 10 m/z isolation window, 1 m/z overlap, collision energy 28, and 15,000 resolution. Files (.raw format) were processed with DIA-NN v2.2.0, analyzing all runs jointly to enable retention-time alignment and match-between-runs; in silico digestion used trypsin-P specificity with one missed cleavage, carbamidomethylation of cysteine as a fixed modification and oxidation of methionine as a variable modification (one per peptide), and N-terminal methionine excision enabled. Protein reannotation used a reviewed *Rattus norvegicus* UniProt reference proteome (accessed March 2024) supplemented with iRT calibration peptides, reversed decoys, and common contaminants, with peptide- and protein-level false discovery rates controlled at 1%. DIA-NN returned 5,002 protein groups across the 24 samples (four groups — CTL, MitoTx, Phe-only, Phe+MitoTx — at six replicates each, with selected technical re-injections per the study metadata). Keratin and bovine fetal-calf-serum contaminants were removed per the recommendations of Frankenfield et al. (19). A protein was retained if it was quantified in at least 4 of 6 replicates in at least one group. Sample outliers were screened by four independent heuristics, namely per-sample missingness above the Tukey upper fence, a Hampel 3×MAD filter on median log₂ intensity, PC1–PC3 Mahalanobis distance (χ² tail, df = 3), and median inter-sample correlation below median − 3×MAD; a sample was removed only when at least three of the four criteria agreed, and no sample met this threshold. Filtering yielded 4,806 proteins × 24 samples, six per group (S1 Table). Abundances were normalized by cyclic-loess in the proteoDA R package (v2.0.0) (20).

Missingness in the normalized matrix was modest (4.7%) and left-censored, where a complete data matrix was required (PCA, clustering) values were imputed with missForest (v1.6.1) (ntree = 100, maxiter = 10) (21) which tracked the non-imputed effect sizes most closely (Spearman 0.98–0.99) without manufacturing differential calls. Differential abundance was reported from the non-imputed matrix — as limma fits each protein independently using only available observations — with limma empirical-Bayes moderation (22), a group-means design, and a duplicateCorrelation replicate block retained at its low consensus correlation (ρ ≈ 0.02) (23). Protein-level significance was determined using the Π-score, a composite metric integrating statistical significance and fold-change magnitude (Π = P.Value^|log₂FC| < 0.05) (24), as the primary threshold; nominal p < 0.05 as well as a Benjamini–Hochberg false discovery rate < 0.10 were also computed as complementary thresholds. Protein set enrichment analysis used ranked-list enrichment (fgseaMultilevel, fgsea v1.38.0) on proteins ordered by the limma moderated-t statistic against pooled rat gene sets (retrieved with msigdbr v26.1.0) between 10-500 targets (Hallmark, Reactome, KEGG, GO-Slim, MitoCarta3.0) with disease, cancer, and infection-related terms excluded. Benjamini–Hochberg correction was applied per database (pathway FDR < 0.05) and redundancy collapsed within- then cross-database by the EnrichmentMap combined overlap/Jaccard coefficient (25). Analyses were performed in R (v4.6.0); PERMANOVA and dispersion tests used vegan (v2.7.3).

Co-expression modules were built with Weighted Gene Co-expression Network Analysis (WGCNA v1.74) (29) on the same 4,806 × 24 matrix using a signed network and signed topological overlap, biweight midcorrelation (maxPOutliers = 0.05) for robustness to outlying samples, and a soft-thresholding power of 8, selected as the lowest power reaching a scale-free fit of R² ≥ 0.85. Blockwise module detection used a minimum module size of 20, deepSplit = 2 and a merge cut height of 0.25, yielding 15 modules plus the unassigned grey set. Modules were first screened contrast-agnostically by an omnibus moderated F test on module eigengenes across the four groups (limma, paired Replicate block), retaining modules at Benjamini–Hochberg FDR < 0.05. Response to each individual contrast was then tested with fry, a self-contained rotation test that asks only whether a module’s own member proteins move. camera and the competitive fgsea adjusted p were computed alongside but are reported as descriptive direction and magnitude only, because a module spanning up to a fifth of the quantified proteome violates the competitive null. fry FDR < 0.05 is the significance gate throughout. Module over-representation used fgsea::fora with the same per-database Benjamini–Hochberg correction and EnrichmentMap deduplication applied to the ranked analysis (32). Per-contrast eigengene shifts from the same limma fit provide the post-hoc comparisons shown as brackets on the module figure. Network construction and module-preservation diagnostics are provided in supplemental data S3 Figure. All stochastic steps used set.seed.

Six complementary clustering pilots interrogated the proteome at the trajectory level (presented in the Results sections). Three fuzzy c-means pilots applied identical e1071::cmeans on gene-level condition means (26), differing only by significance gate (nominal p < 0.05, Π < 0.05, or BH-FDR < 0.10 in ≥1 core contrast). The fuzzifier m was set by the Schwämmle–Jensen estimator (27), and the number of clusters c was chosen from the minimum-centroid-distance elbow (drop < 10% of curve range), capped at ⌊√(N/2)⌋ per the standard cluster-count upper bound (28). A fourth pilot grouped proteins by Weighted Gene Co-expression Network Analysis modules (29) from the pre-built signed-Pearson network and correlated each module eigengene with binary indicator vectors for Disease (Phe-only − CTL), Transplant (MitoTx − CTL), and Rescue (Phe+MitoTx – Phe-only); modules with opposite-sign Disease and Rescue r at |r| ≥ 0.25 were called “Reversal”, a sub-significance floor adopted because n = 24 makes the conventional |r| ≥ 0.40 cutoff collapse every module to “Other” (treated throughout as a ranking aid rather than a per-module inferential claim). A fifth pilot clustered each protein’s four-dimensional log₂FC vector (CTL vs Phe-only, Phe-only vs Phe+MitoTx, CTL vs MitoTx, Interaction) by k-means (nstart = 50, iter.max = 100), with k selected by the gap statistic firstSEmax rule (30) and each cluster labeled by the (Disease, Rescue) sign quadrant of its centroid (Reversed Up, Reversed Down, Concordant Up, Concordant Down, Neutral). A sixth pilot ran Rank-Rank Hypergeometric Overlap (RRHO2; Cahill et al. (31)) between the Disease and Rescue contrasts ranked by signed limma moderated-t (chosen for variance stabilization at n = 24 and consistency with the fgsea cache), tagging genes by quadrant (concordant-up UU, concordant-down DD, reversed UD and DU); for sparse quadrants where the RRHO2 peak-overlap set was < 5 genes, a top-20% percentile fallback (intersection of the top 20% of both ranked lists in the quadrant’s implied direction) was substituted and flagged in both the workbook and figure subtitle. Per-cluster, per-module, and per-quadrant over-representation were tested with fgsea::fora against Hallmark and MitoCarta3.0 in parallel per the two-source pairing guidance of Reimand et al. (32), with the same per-database BH correction. All stochastic steps used set.seed.

#### Statistical Analysis

All results are reported as mean ± standard deviation values and expressed relative to control values. Non-proteomic comparisons were made using one-way ANOVAs, followed by Tukey’s post hoc testing as appropriate. Welch’s t-tests were used to analyze high-resolution respirometry data. Significance was established at p<0.05 throughout except for proteomics data as outlined above. GraphPad Prism 10.6.1 (GraphPad San Diego, CA, USA) was used for all analyses except proteomic data, as discussed above. All supplemental data are available on an online data repository at DOI 10.5281/zenodo.21679814.

## RESULTS

### AIM 1 efficacy of mitochondrial transplant into H9C2 cardiomyocytes

Coincubation of fluorescently labelled L6 myotube mitochondrial isolates with H9C2 cardiomyocytes led to mitochondrial internalization in H9C2 cardiomyocytes (Figure 1A). An increase in certain oxygen consumption rate (OCR) kinetics was observed with mitochondrial transplantation (Figure 1B). Specifically, compared with PBS-treated controls, there was a statistically significant increase in both Complex I, State 3 respiration (p=0.005) and maximal respiration (ET Capacity) (p=0.022) in cardiomyocytes subjected to mitochondrial transplantation. Although Complex IV, State 3 (Complex IV Linked Respiration) was improved with mitochondrial transplantation, it did not reach statistical significance compared with control (p=0.071). Mitochondrial outer membrane integrity was verified utilizing the cytochrome c test (*j_c_*). The control average was 8.6%, and the mitochondrial transplantation average was 6.0% (p=0.525 between conditions) with both being below the 15% quality control threshold.

**FIGURE 1.**
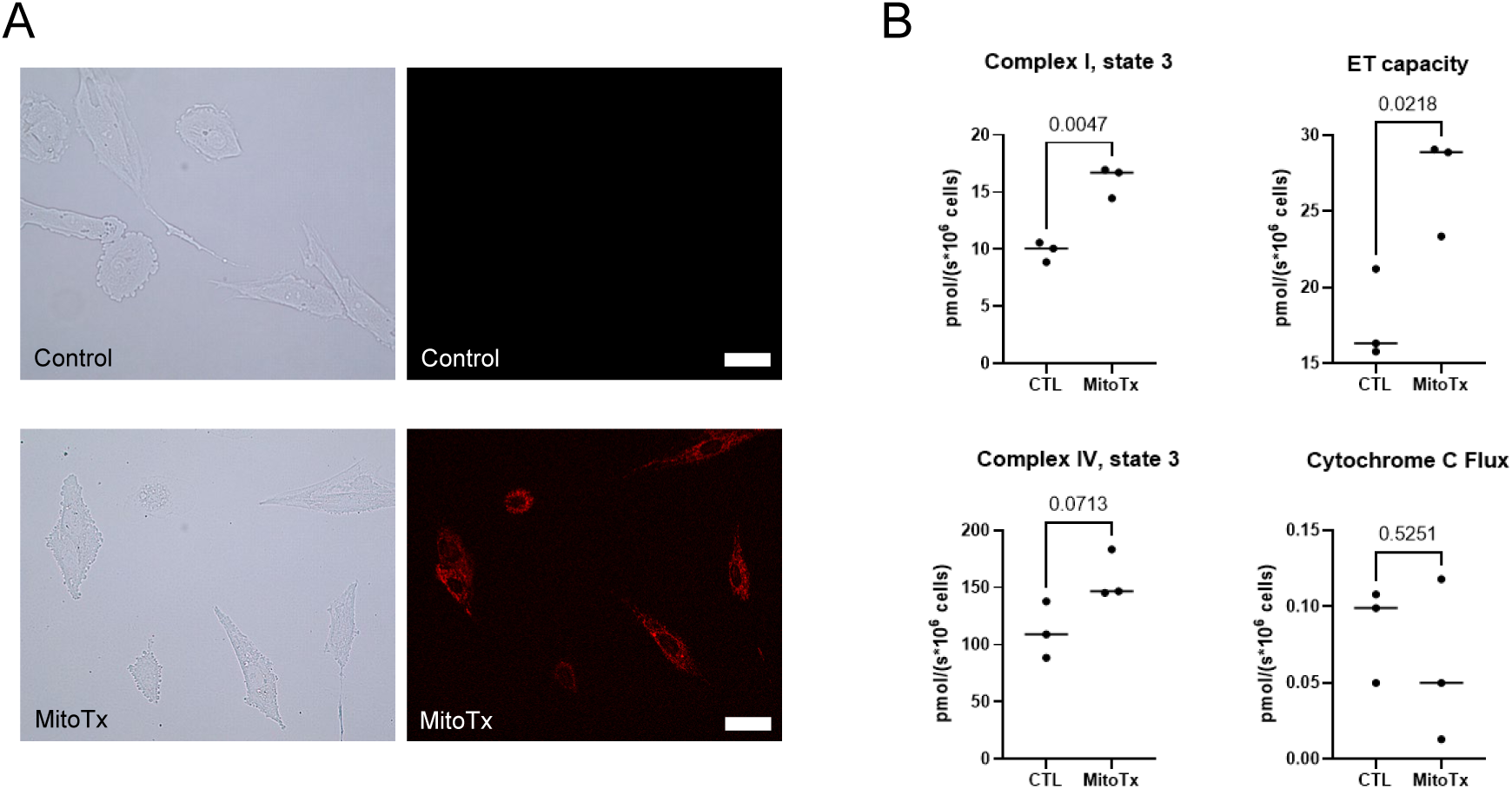
AIM 1 results demonstrating successful L6 mitochondrial transplantation in H9C2 cardiomyoctes. A) Internalization of exogenous L6 myotube mitochondria (pre-labelled with Mito-Tracker Red FM) into non-perturbed H9C2 cardiomyocytes. Control cells received a vehicle treatment of PBS. Fluorescent microscopy revealed perinuclear localization of the stained exogenous mitochondria following coincubation. The brightfield and associated fluorometric images were taken after 24 hours of coincubation using 10x magnification (white bar = 100 µM). B) Respirometry data on n=3 replicates per condition indicates that mitochondrial transplantation increased complex I state 3 (complex I-linked OXPHOS) and maximum capacity of the electron transfer system (ET capacity) without affecting cytochrome c flux. Values are normalized to cell count. Abbreviations: CTL, control cells with no mitochondrial transplantation; MitoTx, cardiomyocytes were coincubated with L6 myotube mitochondrial isolates for 24 hours.

### Phenylephrine dose response test in H9C2 cardiomyocytes prior to AIM 2 experiments

Exposure to 100 μM phenylephrine for 48 hours significantly increased cardiomyocyte size (p=0.031) compared with untreated controls (Figure 2A/B). Note, cardiomyocyte size was numerically larger after exposure to 25 μM and 50 μM phenylephrine compared with control, though these differences were not statistically significant relative to the control condition. The bioenergetics of cardiomyocytes exposed to 100 μM of phenylephrine were compared to those of untreated control cardiomyocytes (Figure 2C). Compared with control, there was no significant difference in Complex I, State 3 respiration (Complex I Linked OXPHOS Capacity) (p=0.70), maximal respiration (ET Capacity) (p=0.99), and Complex IV, State 3 (Complex IV Linked Respiration) (p=0.54) after exposure to 100 μM phenylephrine. Again, mitochondrial outer membrane integrity was verified utilizing the cytochrome c test (*j_c_*). The control average was 5.6%, and the PE average was 6.8%, both below the 15% quality control threshold.

**FIGURE 2.**
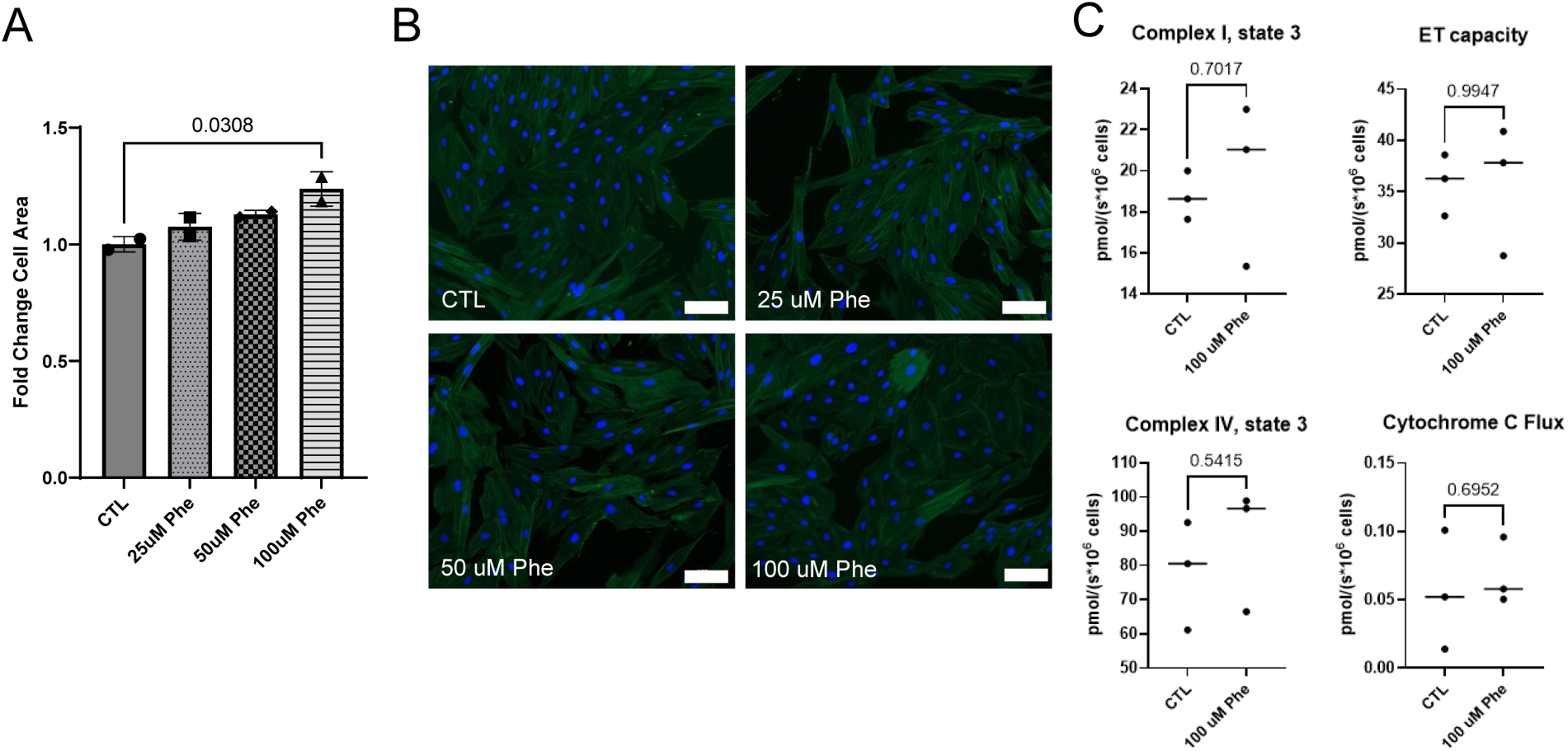
Phenylephrine pilot experiment to ensure cardiomyocyte hypertrophy. A) Cell size differences presented as means ± standard deviation values between Control and different phenylephrine doses (expressed as fold-change from control treatments). Two replicate wells were used for each group, and 5 images were taken randomly per well. ∼20-30 cardiomyocytes were analyzed for morphology per image under 10x magnification (white bar = 100 µM), B) Representative 10x image of each treatment condition. C) Respirometry data on n=3 replicates per condition indicates that 100 μM of phenylephrine did not affect complex I state 3 (complex I-linked OXPHOS), complex IV state 3 (complex IV-linked OXPHOS), maximum capacity of the electron transfer system (ET capacity), or cytochrome c flux. Values are normalized to cell count. Abbreviations: CTL, control cells with no phenylephrine; Phe, 25-100 µM phenylephrine was applied to cells for 24 hours.

### AIM 2 assessments of L6 myotube mitochondrial transplant effects on H9C2 cardiomyocytes treated with phenylephrine

The 100 μM phenylephrine dose, despite not affecting mitochondrial bioenergetics (Figure 2C), was used for mitochondrial transplantation experiments given that it induced cardiomyocyte hypertrophy. Though 100 μM Phe significantly increased cardiomyocyte size compared with untreated control cardiomyocytes, L6 mitochondrial transplantation significantly reduced cardiomyocyte size in phenylephrine-treated H9C2 cells and restored cell size to a level comparable to that of control cells (Figure 3A/B). In a separate set of cells, L6 mitochondria tagged with MitoTracker Red FM were co-incubated with H9C2 cardiomyocytes after the first 24 hours of phenylephrine exposure to ensure that transplantation was not affected by Phe treatments. It was confirmed that mitochondrial internalization occurred in the presence of 100 μM Phe (Figure 3C).

**FIGURE 3.**
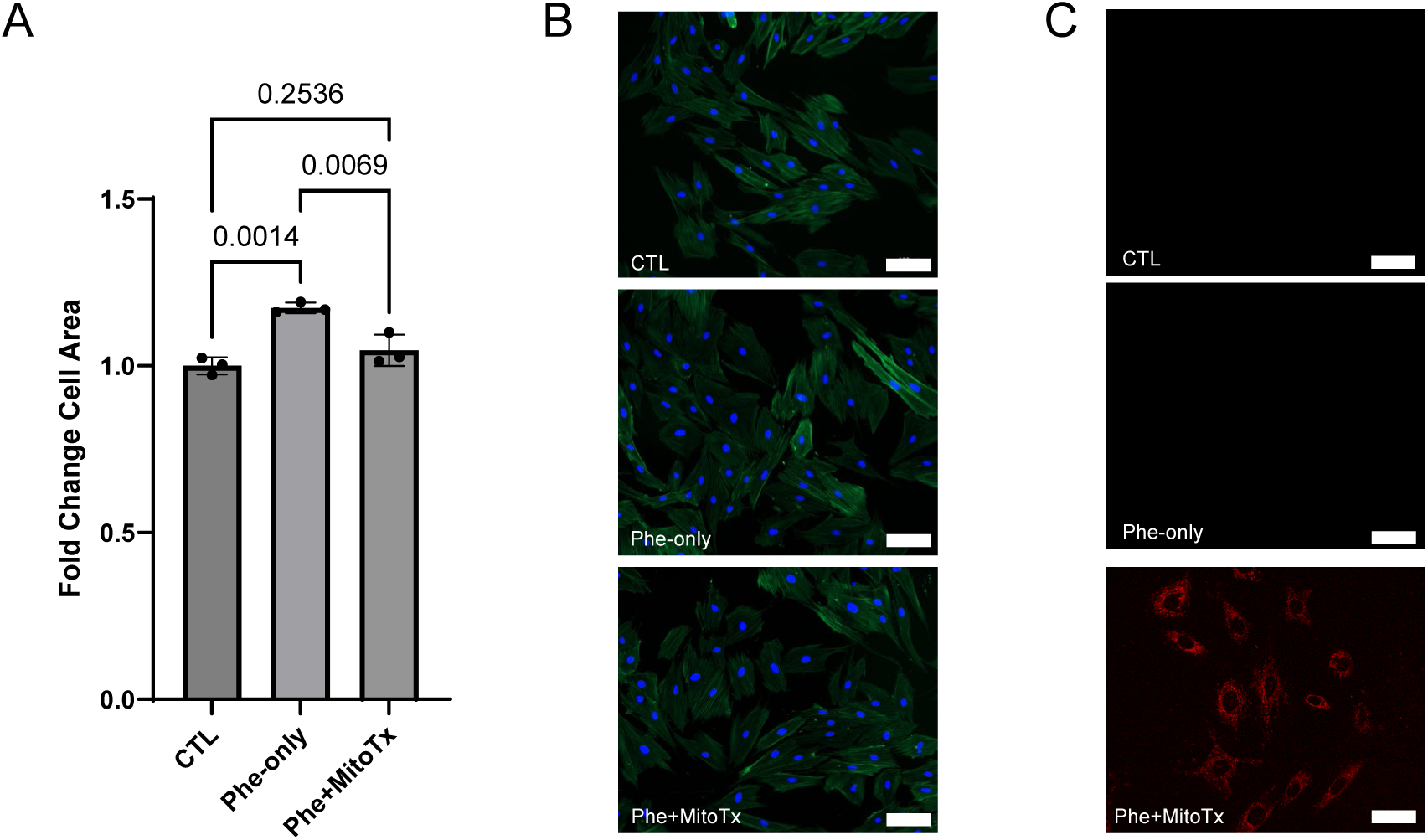
AIM 2 results examining the effects of mitochondrial transplantation on cell size. A) Cell size differences presented as means ± standard deviation values between Control, Phe-only, and Phe+MitoTx (expressed as fold-change from control treatments). Three replicate wells were used for each group and ∼20-30 cardiomyocytes per treatment were analyzed for morphology per image under 10x magnification (white bar = 100 µM). B) Phalloidin+DAPI representative 10x images of Control, Phe-only, and Phe+MitoTx. C) Bottom micrograph demonstrates internalization of exogenous L6 myotube mitochondria (pre-labelled with Mito-Tracker Red FM) into H9C2 cardiomyocytes following coincubation with 100 μM Phe relative to the other two images whereby mitochondrial transplantation was not applied. Abbreviations: CTL, control cells with no phenylephrine and no mitochondrial transplantation; Phe-only, 100 µM phenylephrine was applied to cells for 48 hours; Phe+MitoTx, cardiomyocytes were treated with 100 µM phenylephrine for 48 hours and transplanted with L6 myotube mitochondrial isolates for the last 24 of the 48 hours.

### AIM 2 proteomic and bioinformatics results

Given that mitochondrial transplant reduced Phe-induced cardiomyocyte hypertrophy, we were interested in leveraging proteomics and bioinformatics to examine molecular phenotype shifts between the AIM 2 treatments. As outlined in the methods, these experiments differed from Figure 3 experiments given that a fourth group (MitoTx-only) was also included to parse out Phe-, MitoTx, and interaction effects. Figure 4 summarizes the global proteome response. Principal-component analysis resolved the four groups across two axes (PC1 23.8%, PC2 12.2%; Figure 4A). Group structure was significant (PERMANOVA R² = 0.219, p < 0.001) with homogeneous dispersion (betadisper p = 0.38). The intervention axis dominated the pairwise tests whereby L6 mitochondrial transplantation (CTL vs. MitoTx R² = 0.183, p = 0.006, BH q = 0.008) and rescue (Phe-only vs. Phe+MitoTx R² = 0.165, p = 0.004, q = 0.008) separated cleanly whereas the disease axis did not move global structure (CTL vs. Phe-only R² = 0.095, p = 0.390, q = 0.390). Sample coordinates, PERMANOVA output and the supporting ordination variants are given in supplemental data S1 Table and S1 Figure.

**Figure 4.**
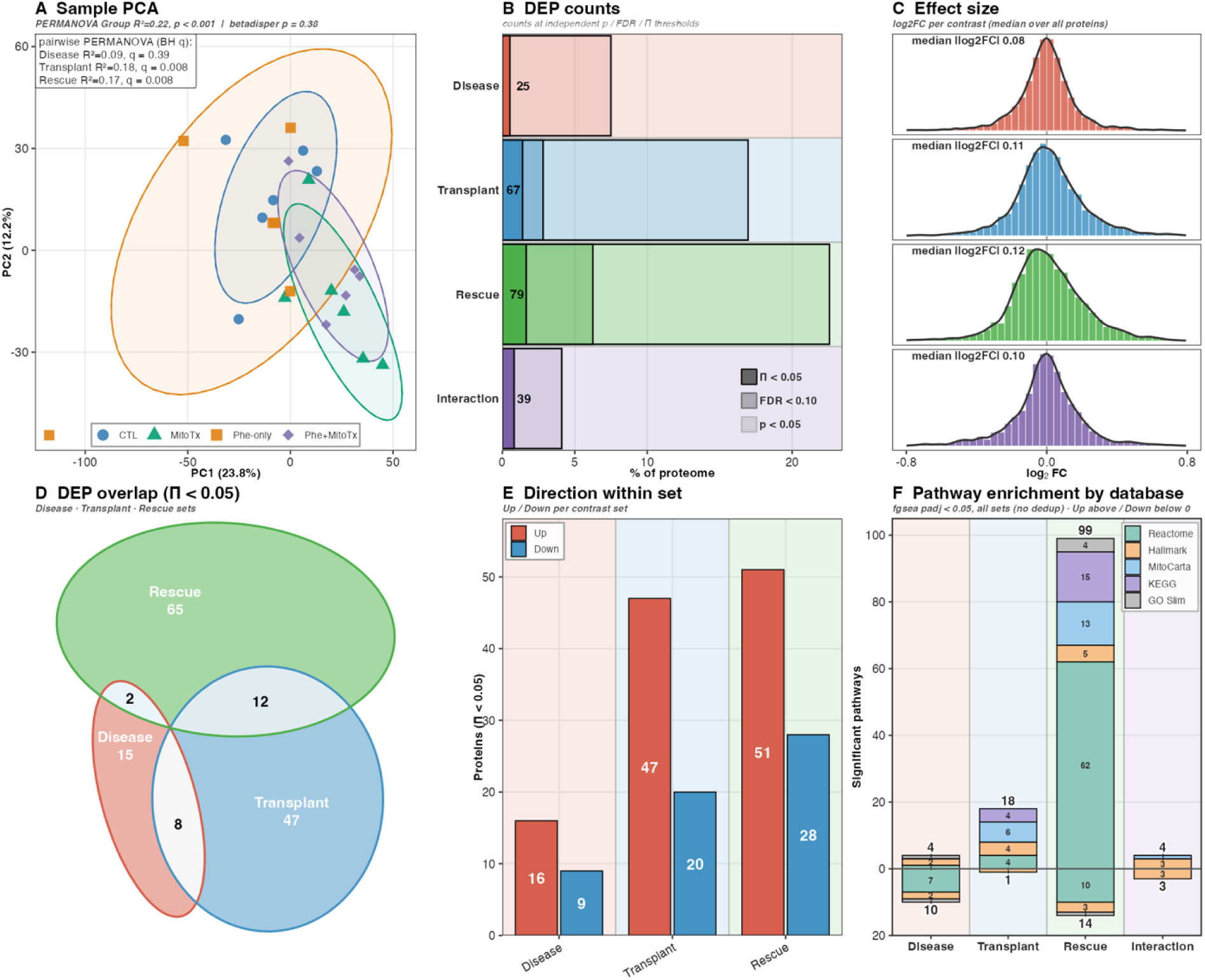
AIM 2 proteomic contrasts between treatments. A) PCA on the missForest-imputed matrix (24 samples (n=6 per treatment) x 4,806 proteins, 80% confidence ellipses; PC1 23.8%, PC2 12.2%). B) Differentially abundant proteins per contrast as % of proteome, split Up versus Down, at three cutoffs (p < 0.05, BH-FDR < 0.10, Π < 0.05); Π < 0.05 is reported throughout. C) log2FC distribution per contrast. D) Area-proportional Euler diagram of Π < 0.05 proteins across Disease, Transplant, and Rescue. E) Up/Down direction strip for the same proteins. F) Significant-pathway counts per contrast and direction before deduplication (five-database, fgsea BH-FDR < 0.05); deduplicated representative counts are given in Fig. 5 and supplemental data S2 Table. Abbreviations: CTL, control cells with no phenylephrine and no mitochondrial transplantation; Phe-only, 100 µM phenylephrine was applied to cells for 48 hours; MitoTx, cardiomyocytes were transplanted with L6 myotube mitochondrial isolates for the 24 hours; Phe+MitoTx, cardiomyocytes were treated with 100 µM phenylephrine for 48 hours and transplanted with L6 myotube mitochondrial isolates for the last 24 of the 48 hours.

Figure 4B displays proteins altered between contrasts according to significance thresholding (nominal p < 0.05, BH-FDR < 0.10, and Π < 0.05), and Figure 4C displays log_2_FC per contrast. Using Π < 0.05 as our exploratory significance threshold yielded 149 unique differentially abundant proteins across the Disease, Transplant and Rescue contrasts (Figure 4D). There was no three-way intersection and only two proteins shared between disease and rescue, both of which reversed direction (Abhd10, Pcdh1); hence, the reversal is read at the directional and pathway level, not single-protein overlap (Figure 4D). Disease and Rescue share the Phe-only group and are therefore anticorrelated at r = −0.50 by construction. A permutation null that shuffles group labels within replicate reproduced the observed proteome-wide anticorrelation almost exactly (observed r = −0.538 against a null median of −0.503, p = 0.45), so the apparent opposition between these two contrasts carried no evidence of reversal. The comparison this design does support Transplant against Rescue, which share no group and are orthogonal: these two contrasts agree proteome-wide at r = 0.797 against a null centered near zero (null median −0.035, p < 0.003), indicating that the L6 transplantation drove one program whether or not the cell is under phenylephrine stress (supplemental data S4 Figure). The rescue contrast carried the largest signature (79 proteins; 51 up, 28 down), exceeding transplantation (67; 47/20), interaction (39; 12/27), and the focal disease signature (25; 16/9) (Figure 4E). Pathway enrichment followed the same ordering. At fgsea BH-FDR < 0.05 the rescue contrast returned 113 significant sets, against 19 for transplantation, 14 for disease and 7 for interaction (Fig. 4F). After EnrichmentMap deduplication these collapsed to 51, 14, 10 and 7 representative programs respectively, leaving rescue by far the richest response (Fig. 5, supplemental data S2 Table).

**Figure 5.**
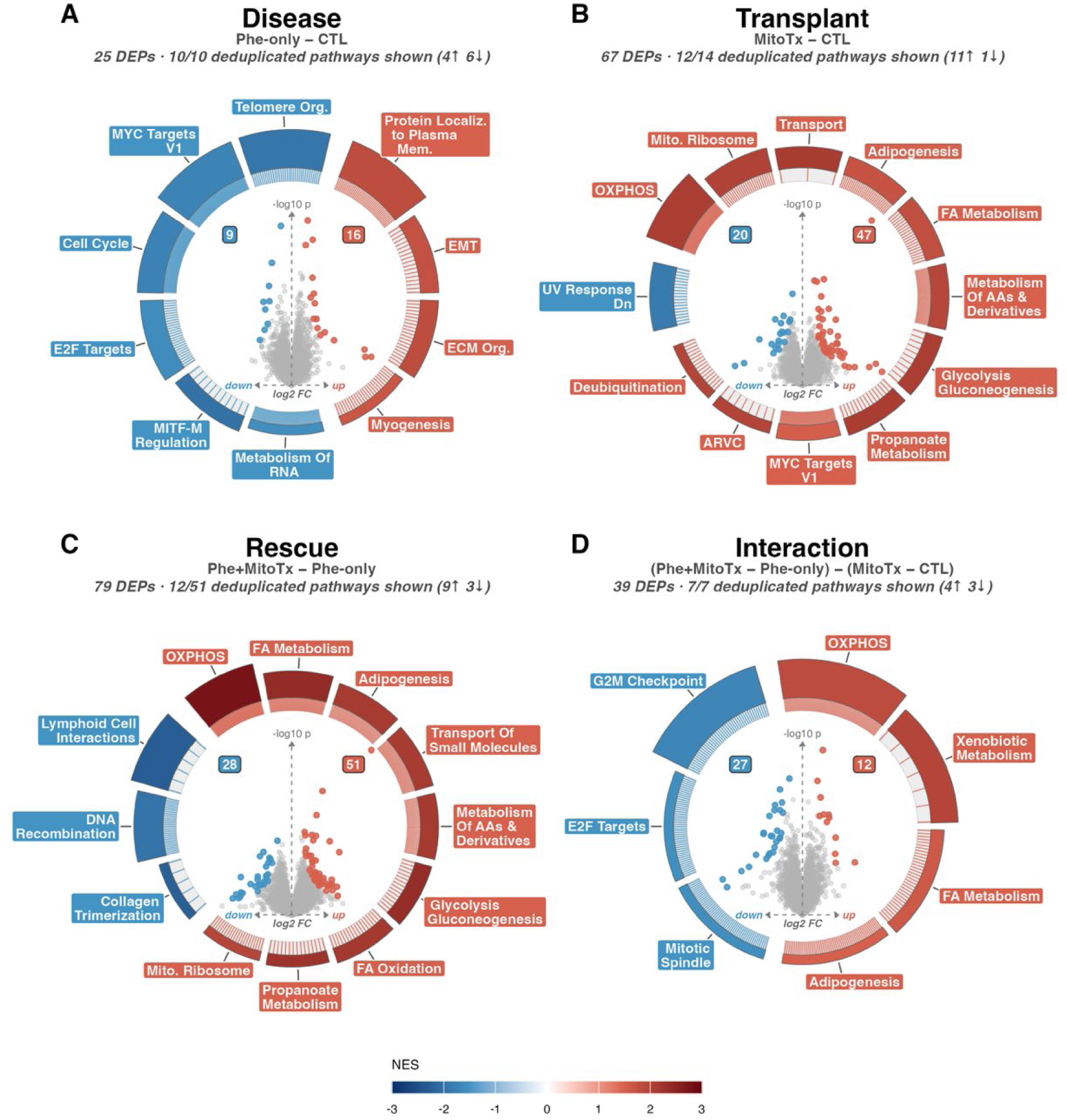
AIM 2 pathway analysis. Per-contrast enrichment volcano-in-ring panels for: A) Disease contrast (Phe-only vs. CTL), B) Transplant contrast (MitoTx vs. CTL), C) Rescue contrast (Phe+MitoTx vs. Phe-only), and D) Interaction contrast ((Phe+MitoTx – MitoTx) – (Phe-only – CTL)). Note that each panel pairs a central per protein volcano (log2FC against significance, with DEP counts) with a surrounding ring of pathways drawn from a pooled five-database lens (Hallmark, Reactome, KEGG, GO-Slim, MitoCarta) at fgsea BH-FDR < 0.05 and collapsed by EnrichmentMap overlap/Jaccard deduplication. Each panel subtitle gives the number of representative programs displayed out of the total surviving deduplication (Disease 10/10, Transplant 12/14, Rescue 12/51, Interaction 7/7); arc height is −log10 FDR and arc colour is NES. Abbreviations: CTL, control cells with no phenylephrine and no mitochondrial transplantation; Phe-only, 100 µM phenylephrine was applied to cells for 48 hours; MitoTx, cardiomyocytes were transplanted with L6 myotube mitochondrial isolates for 24 hours; Phe+MitoTx, cardiomyocytes were treated with 100 µM phenylephrine for 48 hours and transplanted with L6 myotube mitochondrial isolates for the last 24 of the 48 hours.

Figure 5 resolves the pathway program unique to each contrast. The Disease contrast (Phe-only vs. CTL) returned 10 representative pathways after deduplication, all of which are displayed (4 up, 6 down), with an upregulation in extracellular-matrix/de-differentiation arm (ECM organization, epithelial-mesenchymal transition, myogenesis) and down-regulation in proteins involved in proliferation (MYC and E2F targets, cell cycle) (Figure 5A). The Transplant contrast (MitoTx vs. CTL) returned 14 pathways, of which the top 12 most significant are displayed (11 up, 1 down), most of which were related to mitochondrial and metabolic adaptation (OXPHOS, mitoribosome, mitochondrial transport, fatty-acid and amino-acid metabolism; full ranked list in supplemental data S2 Table) (Figure 5B). The Rescue contrast (Phe+MitoTx vs. Phe-only) returned 51 pathways, the top 12 are displayed (9 up, 3 down), and these pathways possessed similarities to the Transplant contrast. However, oxidative phosphorylation was the strongest pathway revealed in this contrast when examining all contrasts (NES = 2.76, p = 3.5×10⁻¹⁸, BH-FDR = 1.7×10⁻¹⁶) (Figure 5C). The Interaction contrast ((Phe+MitoTx – MitoTx) – (Phe-only – CTL)) returned 7 pathways (4 up, 3 down), and again pathways related to mitochondrial and metabolic adaptation were upregulated (Figure 5D). Per-contrast enrichment, the transplantation’s disease-specificity, and the disease-selected sets read across contrasts are shown in supplemental data S2 Figure.

Protein-level contrasts treat each protein independently. Hence, as a final analysis we were also interested in whether transplantation moved coordinated groups of proteins. Weighted gene co-expression network analysis on the same 4,806 × 24 matrix resolved 15 modules plus an unassigned grey set, ranging from 46 to 1,091 proteins. Screening modules contrast-agnostically by an omnibus moderated F test across the four groups identified five responsive modules at FDR < 0.05: blue (801 proteins, q = 0.011), greenyellow (110, q = 0.011), turquoise (1,091, q = 0.014), brown (421, q = 0.028) and tan (83, q = 0.031). Over-representation assigned turquoise to mitochondrial energetics (oxidative phosphorylation, mitochondrial matrix, fatty-acid and organic-acid metabolism) and blue to adhesion and matrix biology (cadherin and actin binding, endoplasmic-reticulum lumen, epithelial-mesenchymal transition, extracellular-matrix organization). Greenyellow carried a single significant term, pyruvate metabolism, and brown returned none, leaving it responsive but functionally unannotated (Fig. 6, supplemental data S3 Table).

**Figure 6.**
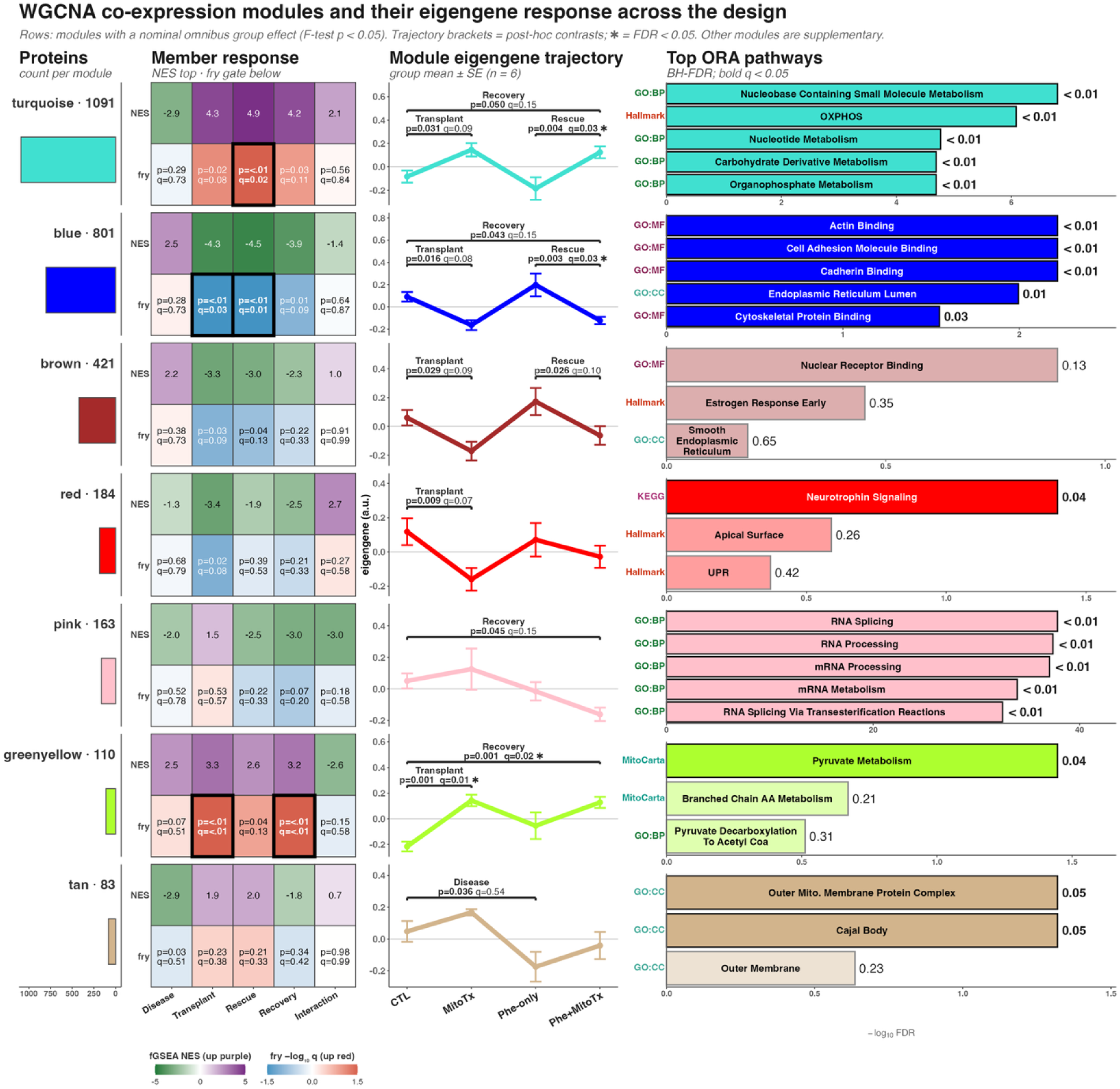
AIM 2 co-expression module analysis. Co-expression modules responsive to treatment. A) Module size in proteins. B) Module eigengene across the four treatment groups (group mean ± SE, n = 6); the omnibus moderated F FDR is printed per module and starred at FDR < 0.05. C) Top five over-representation terms per module (per-database BH-FDR; bars at full color are FDR < 0.05, washed-out bars are not significant). D) Module normalized enrichment score across the Disease, Transplant, Rescue and Interaction contrasts, shown as descriptive direction and magnitude only; a ring outline marks cells clearing the fry self-contained rotation gate at FDR < 0.05. The full module × contrast fry grid, including the Recovery and disease-after-transplant contrasts, is in supplemental data S3 Table. Abbreviations: CTL, control cells with no phenylephrine and no mitochondrial transplantation; MitoTx, cardiomyocytes transplanted with L6 myotube mitochondrial isolates for 24 hours; Phe-only, 100 µM phenylephrine applied to cells for 48 hours; Phe+MitoTx, cardiomyocytes treated with 100 µM phenylephrine for 48 hours and transplanted with L6 myotube mitochondrial isolates for the last 24 of the 48 hours; NES, normalized enrichment score.

Testing every module against every contrast with fry, a self-contained rotation test, 5 of 90 module × contrast cells cleared FDR < 0.05: greenyellow on Transplant (q = 0.005) and on Recovery (q = 0.008), turquoise on Rescue (q = 0.017), and blue on Rescue (q = 0.011) and on Transplant (q = 0.031). Turquoise and greenyellow moved up and blue moved down in each of these cells, so the coordinated response to the transplant was a rise in mitochondrial-energetic modules alongside a fall in the adhesion and matrix module. No module cleared this gate on Disease, on Disease-after-transplant or on the Interaction. The competitive statistics disagree sharply and are not read as support: camera returns FDR < 0.05 for 70 of the same 90 cells, with adjusted p values as small as 1×10⁻⁷¹ for turquoise, and module-level fgsea is similarly extreme (BH-adjusted p to 2×10⁻¹⁰) because a module spanning a fifth of the quantified proteome breaks the competitive null. Module enrichment scores are therefore reported as direction and magnitude only. At six replicates per group these module tests remain exploratory, and the confirmatory signal in this study is the protein-level differential abundance reported above.

## DISCUSSION

We sought to examine the effects of exogenous mitochondrial transplantation in phenylephrine-treated cardiomyocytes. AIM 1 provides relatively straightforward methods for *in vitro* mitochondrial transplantation that are conceivably replicable across laboratories. AIM 2 revealed that Phe+MitoTx blunted the hypertrophic cardiomyocyte response (versus Phe-only treatments) while MitoTx (regardless of drug presence) upregulated a proteomic signature related to mitochondrial and metabolic adaptations. The study design supports this equivalence directly. The transplant-in-healthy contrast (MitoTx − CTL) and the rescue contrast (Phe+MitoTx − Phe-only) share no experimental group, so they are statistically independent, yet their protein changes agree across the proteome (r = 0.797, against a permutation null centered near zero). The broader implications of these findings are presented in the remainder of the discussion.

The most notable finding herein was the favorable effect of L6 myotube mitochondrial transplant on phenylephrine-induced cardiomyocyte hypertrophy. Several studies have reported phenylephrine’s hypertrophic effects on H9C2 cardiomyocytes (33–35), and our data are in line with these prior studies. However, it is notable that the phenylephrine insult produced a focal rather than global perturbation given that our treatment protocol (100 µM for 48 hours): i) did not alter the bulk proteome, as only 25 proteins were altered at Π < 0.05, and ii) did not disrupt mitochondrial respiration. Notwithstanding, the proteome enrichment profile contrasts revealed that proteins related to matrix deposition and epithelial-mesenchymal/de-differentiation were significantly altered with phenylephrine treatments, and prior transcriptomics work suggests these signatures coincide with phenylephrine-induced cardiac hypertrophy in rodents (36).

Given that our *in vitro* model was established to provide potential direction for *in vivo* studies, a comparison of our data with two notable studies in this area is provided. Weixler and colleagues performed a two-part study with an *in vitro* arm examining the effects of mitochondrial transplant on angiotensin II-induced cardiomyocyte hypertrophy along with an *in vivo* arm in pigs whereby RV failure was surgically induced (15). These authors reported that mitochondrial transplantation restored ATP levels in hypertrophied cardiomyocytes to control levels, with skeletal muscle-derived mitochondria (both fast- and slow-twitch muscles) proving equally effective as cardiac muscle-derived mitochondria in reversing the bioenergetic deficits. Ali Pour et al. (37) studied the bioenergetic consequences of exogenous mitochondrial transplantation from L6 skeletal muscle cells into unperturbed H9C2 cardiomyocytes. There was acute enhancement in bioenergetics two days after mitochondrial transplantation, but this effect was diminished following longer-term assessments (after 7 days). Based on the long-term bioenergetics profile at 28 days, there were no negative bioenergetic consequences related to mitochondrial transplantation. Collectively, these two studies along with our current study indicate that exogenous mitochondrial transplantation, particularly from skeletal muscle-derived sources, can restore bioenergetic function in stressed cardiomyocytes. It is also notable that the collective evidence indicates that mitochondrial transplantation may attenuate hypertrophic remodeling. However, given the limited preclinical data in this area, the durability of these effects and their translation to intact cardiac tissue warrants further investigation.

### Limitations

*In vitro* models are valuable for isolating specific biological mechanisms, but they inherently fail to replicate the complex, dynamic environment of living organisms, especially in clinical contexts such as RV failure. Because of this significant limitation, neither this nor previously published preclinical studies in this area can be used to predict the clinical efficacy or safety of mitochondrial transplantation for treatment of conditions such as RV failure in humans. We also acknowledge that this was an exploratory study using phenylephrine-induced cardiomyocyte hypertrophy. In our model, phenylephrine produced a small but focused disease signature.

Morphologically, phenylephrine significantly increased cell size. This aligned with our pathway enrichment analysis of the proteomic dataset, which showed upregulation of hypertrophic mechanisms and suppression of proliferation programming. However, we did not observe that phenylephrine affected mitochondrial respiration via high-resolution respirometry. We also did not measure ROS in our experiments to see if phenylephrine affected parameters other than oxygen consumption rate. Future studies should investigate if mitochondrial transplantation affects phenotypic and/or molecular outcomes in cardiomyocytes treated with other drugs or agents that serve to provide different stress responses (e.g., hydrogen peroxide or hypoxia/reoxygenation to simulate ischemic and oxidative stress, and high glucose or palmitate to recapitulate metabolic cardiomyopathy). Two limitations apply to the proteomic analysis.

First, the six pairwise contrasts were not independent (i.e., contrasts that share a group, such as disease and rescue, are correlated by arithmetic alone), so we do not read their opposition as biological reversal. Second, at six replicates per group the co-expression module tests are exploratory, and the confirmatory signal remains the protein-level differential abundance which was weak at FDR significance thresholds.

### Conclusions

In summary, this study demonstrates that exogenous mitochondrial transplantation from L6 myotubes can blunt phenylephrine-induced hypertrophic remodeling while simultaneously upregulating mitochondrial and metabolic proteomic signatures in H9C2 cardiomyocytes. These findings, considered alongside prior preclinical evidence, continue to support mitochondrial transplantation as a bioenergetically favorable and potentially cardioprotective intervention. However, additional cell culture alongside rigorous *in vivo* studies across diverse cardiac stress models are needed before the translational potential of this approach for the treatment of RV failure can be meaningfully evaluated.

## ADDITIONAL INFORMATION

## Acknowledgements

We graciously thank Dr. Gaurav Choudhary (Brown University, USA) and Dr. Joshua Godwin (Penn State University) for insights during these experiments.

## Author contributions

Conceptualization, N.J.K, G.J.K, B.A.R.; funding acquisition, M.D.R.; investigation and methodology, N.J.K, G.J.K., D.T.L., S.C.N., M.D.B., A.I.; formal analysis, N.J.K., G.J.K., A.I.; supervision, A.N.K., D.T.B., D.J.M., C.B.M., M.D.R., writing-original draft, N.J.K., D.T.L., M.D.R.; review and editing, all co-authors; final approval of manuscript, all co-authors.

## Availability of data and materials

Raw data related to the current study outcomes will be provided upon reasonable request by emailing the corresponding author. Processed proteomics data are available at DOI 10.5281/zenodo.21679814. Analysis code is available at https://github.com/Dustyn-T-Lewis/Mito_2026.

## SUPPLEMENTAL APPENDIX

S1 Table. Proteome overview: PCA sample coordinates, PERMANOVA and dispersion tests, differentially abundant protein counts at each threshold, per-contrast differential-abundance tables for all six contrasts, overlap membership across the Disease, Transplant and Rescue sets, and per-database pathway counts.

S2 Table. Pathway enrichment: every tested pathway per contrast with its NES, nominal p, per-database FDR, set size, figure-inclusion flag, and deduplication status and merge target.

S3 Table. Module landscape: module sizes and omnibus F results, eigengene contrast statistics including the Recovery and Disease-after-transplant contrasts, module preservation, eigengene trajectories, per-module over-representation, and top hub proteins by module membership.

S1 Figure. Ordination and enrichment diagnostics for the proteome overview.

S2 Figure. Per-contrast enrichment and the transplantation’s disease-specificity.

S3 Figure. Module construction, preservation and hub structure.

S4 Figure. Contrast-coupling permutation nulls.

